# The genome of the soybean cyst nematode (*Heterodera glycines*) reveals complex patterns of duplications involved in the evolution of parasitism genes

**DOI:** 10.1101/391276

**Authors:** Rick Masonbrink, Tom R. Maier, Usha Muppiral, Arun S. Seetharam, Etienne Lord, Parijat S. Juvale, Jeremy Schmutz, Nathan T. Johnson, Dmitry Korkin, Melissa G. Mitchum, Benjamin Mimee, Sebastian Eves-van den Akker, Matthew Hudson, Andrew J. Severin, Thomas J. Baum

## Abstract

*Heterodera glycines*, commonly referred to as the soybean cyst nematode (SCN), is an obligatory and sedentary plant parasite that causes over a billion-dollar yield loss to soybean production annually. Although there are genetic determinants that render soybean plants resistant to certain nematode genotypes, resistant soybean cultivars are increasingly ineffective because their multi-year usage has selected for virulent *H. glycines* populations. The parasitic success of *H. glycines* relies on the comprehensive re-engineering of an infection site into a syncytium, as well as the long-term suppression of host defense to ensure syncytial viability. At the forefront of these complex molecular interactions are effectors, the proteins secreted by *H. glycines* into host root tissues. The mechanisms of effector acquisition, diversification, and selection need to be understood before effective control strategies can be developed, but the lack of an annotated genome has been a major roadblock. Here, we use PacBio long-read technology to assemble a *H. glycines* genome of 738 contigs into 123Mb with annotations for 29,769 genes. The genome contains significant numbers of repeats (34%), tandem duplicates (18.7Mb), and horizontal gene transfer events (151 genes). Using previously published effector sequences, the newly generated *H. glycines* genome, and comparisons to other nematode genomes, we investigate the evolutionary mechanisms responsible for the emergence and diversification of effector genes.

## Background

The soybean cyst nematode (SCN) *Heterodera glycines* is considered the most damaging pest of soybean and poses a serious threat to a sustainable soybean industry [1]. *H. glycines* management relies on crop rotations, nematode resistant crop varieties, and a panel of biological and chemical seed treatments. However, cyst nematodes withstand adverse conditions and remain dormant for extended periods of time, and therefore, are difficult to control. Furthermore, the overuse of resistant soybean varieties has stimulated the proliferation of virulent nematode populations that can infect these varieties [2]. Hence, there continues to be a strong need to identify, develop, and implement novel sources of nematode resistance and management strategies.

*H. glycines* nematodes are obligate endoparasites of soybean roots. Once they emerge from eggs in the soil, they find nearby soybean roots and penetrate the plant tissue where they migrate in search for a suitable feeding location near the vascular cylinder. The now sedentary *H. glycines* convert adjacent root cells into specialized, fused cells that form the feeding site, termed syncytium [3]. The parasitic success of *H. glycines* depends on the formation and long-term maintenance of the syncytium, which serves as the sole source of nutrition for the remainder of its life cycle. Host finding, root penetration, syncytium induction, and the long-term successful suppression of host defenses are all examples of adaptation to a parasitic lifestyle. At the base of these adaptations lies a group of nematode proteins that are secreted into plant cells to modify host processes [4]. Intense research is focused on identifying these proteins, called effectors, and to elucidate their complex functions. To date, over 80 *H. glycines* effectors have been identified and confirmed [5, 6], although many more remain to be discovered. Characterization of some known effectors has provided critical insights into the parasitic strategies of *H. glycines*. For example, these studies revealed that effectors are involved in a suite of functions, including defense suppression, plant hormone signaling alteration, cytoskeletal modification, and metabolic manipulation (reviewed by [7–10]). However, research has yet to provide a basic understanding of the molecular basis of virulence, i.e., the ability of some nematode populations to infect soybean plants with resistance genes, while other nematode populations are controlled by these resistance genes.

*H. glycines* populations are categorized into Hg types based on their virulence to a panel of soybean cultivars with differing resistance genetics [2, 11]. Based on the Hg type designation, growers can make informed decisions on soybean cultivar choice. To date, the Hg type designation can only be ascertained through time-consuming and expensive greenhouse experiments. However, once the genetic basis for virulence phenotypes has been explored, it is conceivable that molecular tests can be developed to make Hg type identification fast and reliable.

Resistant soybean cultivars are becoming less effective, as *H. glycines* populations alter their Hg type designation as a function of the soybean resistance genes to which the nematode population is exposed. In other words, when challenged with a resistant soybean cultivar for an extended duration, the surviving nematodes of an otherwise largely non-virulent *H. glycines* population will eventually shift towards a new Hg type that is virulent on resistant soybean cultivars [2]. It is unknown if this phenomenon solely relies on the selection of virulent genotypes already present within a given nematode population, or if *H. glycines* wields the power to diversify an existing effector portfolio to quickly infect resistant soybean cultivars. In addition, such genetic shifts appear to be distinct across populations with the same pathotype, indicating populations can independently acquire the ability to overcome host resistance [12]. Understanding these and other questions targeting the molecular basis of *H. glycines* virulence are critical for sustainable soybean production in a time when virulent nematodes are becoming more prevalent.

Scientists can finally start answering such questions, as we are presenting the first complete and fully annotated *H. glycines* genome along with single-nucleotide polymorphisms (SNPs) associated with fifteen *H. glycines* populations of differing virulence phenotypes. PacBio long-reads were assembled and annotated into 738 contigs of 123Mb containing 29,769 genes. The *H. glycines* genome has significant numbers of repeats (34% of the genome), tandem duplications (14.6Mb), and horizontal gene transfer events (151 genes). Using this genome, we explored potential mechanisms for how effectors originate, duplicate, and diversify. Specifically, we found that effectors are frequently associated with tandem duplications, DNA transposons, and LTR retrotransposons. Additionally, we have leveraged RNA-seq data from pre-parasitic and parasitic nematodes and DNA sequencing across 15 *H. glycines* populations to further characterize effector expression and diversity.

## Results

*H. glycines* genomic DNA was extracted, and PacBio sequencing generated 2.4 million subreads with an average length of 7.6kb corresponding to a coverage of 141x at an estimated genome size of 129MB [13]. Due to the high level of heterozygosity of *H. glycines* populations, our early PacBio-only assemblies using Falcon and Falcon-Unzip resulted in an abundance of heterozygous contigs (haplotigs). Therefore, we reduced the heterozygosity of the original reads using a combination of Falcon, CAP3, and manual scaffolding of the assembly graph in Bandage. The final assembly was polished with Quiver and contains 738 contigs with an N50 of 304kb and a total genomic content of 123,846,405 nucleotides (Figure 1). We confirmed the assembly to be free of contamination using Blobtools (4.8.2) (Figure S1) and validated for completeness by alignment of raw data: 88% of the RNA-seq and 93% of the PacBio preads (Table S1). In addition, approximately 72% of the 982 Nematoda-specific BUSCO genes are complete in the *H. glycines* genome, which is comparable to BUSCO scores in other Tylenchida genomes (Table S2). Remarkably, only 56% of the BUSCO genes in *H. glycines* are single-copy, while 16% were duplicated, a statistic that is comparable to the allopolyploid root-knot nematode *Meloidogyne incognita* (Table S2)[14] [15, 16]. A phylogenetic tree (Figure 1) confirming the established phylogeny was generated using 651/982 single-copy BUSCO genes shared by at least three species among *H. glycines, Globodera pallida, Globodera ellingtonae, Globodera rostochiensis, Meloidogyne hapla, M. incognita*, and *Bursaphelenchus xylophilus*.

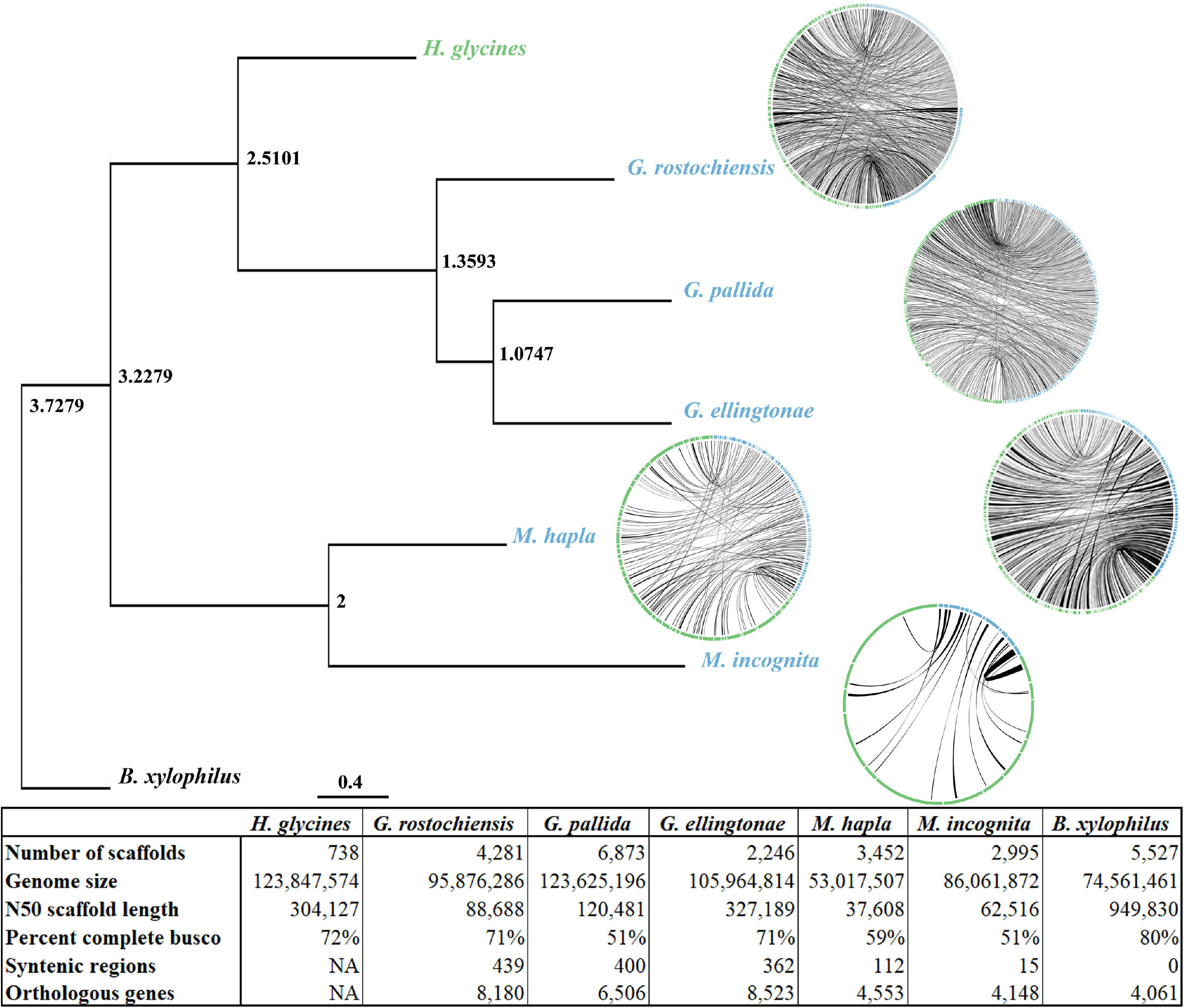

Gene annotations were performed using Braker on an unmasked assembly, as multiple known effector alignments were absent from predicted genes when the genome was masked (Figure S2). While all known effectors are present in the assembly, the resulting gene count of 29,769 also includes a number of expressed repetitive elements (12,357). A wide variety of transcriptional sequencing was used as input for gene annotations, including 230 million RNA-seq reads from both pre-parasitic and parasitic J2 *H. glycines* nematodes, 34,041 iso-seq reads from early, middle and late life stages of both a virulent and an avirulent strain, and the entirety of the *H. glycines* ESTs in NCBI (35,796). In total, 57,893 transcripts from 25,698 genes (2.25 per gene, on average) were annotated, but 26,608 transcripts from 7,063 genes (3.78 per gene, on average) were identified with at least 1 TPM (transcripts per million). Across the various biological groups, 4,114 transcripts from 2,194 genes were attributed with functional consequences to the protein structure as a result of alternative splicing (Table S3). The most abundant alternative splicing events were intron retention (30%) and non-mediated decay (15%), with 70% of alternative splicing events changing open reading frame length (Figure S3 & S4). In comparison, previous work utilizing a de novo transcriptome assembly approach discovered 71,093 genes with 147,910 (2.08 average) transcripts in *H. glycines* [17] thus demonstrating the importance of using an assembled genome as a part of the transcriptomics pipeline [18].

With the genome and gene annotations, we investigated effector genes, which are likely to be involved in virulence. Effector transcript sequences originate in the esophageal glands, which are comprised of two secretory cell types: the subventral and dorsal gland cells. Dorsal gland-expressed genes (DOGs) are mostly active during the later parasitic stages when syncytial development is initiated and progressing. In *Globodera* cyst nematode species, a putative regulatory promoter motif of dorsal gland cell expression, the DOG box, was recently identified [12]. To determine whether the regulation of dorsal gland cell expression in *Heterodera* species may be under similar control, we generated a non-redundant list of putative orthologues of known dorsal gland effectors from cyst nematodes. This included all known dorsal gland effectors, the large family of recently characterized glutathione synthetase-like effectors [19], and all DOG-box associated effectors of *G. rostochiensis*. A total of 128 unique dorsal gland effector-like loci were identified in the genome, their promoter regions were extracted and compared to a random set of non-effector gene promoters using a non-biased differential motif discovery algorithm. Using this approach, a near-identical DOG box motif was identified (Figure 2A), enriched on both strands of dorsal gland effector-like loci promoters approximately 100-150 bp upstream of the start codon (Figure 2B). DOG box motifs occur at a greater frequency in promoters than in the genome, however their presence in a promoter is only a modest prediction of secretion (Figure 2C). Taken together this suggests that the cis-regulatory elements controlling dorsal gland effector expression may be a conserved feature in cyst nematodes, predating at least the divergence of *Globodera* and *Heterodera*, and thus have been conserved for over 30 million years of evolution.

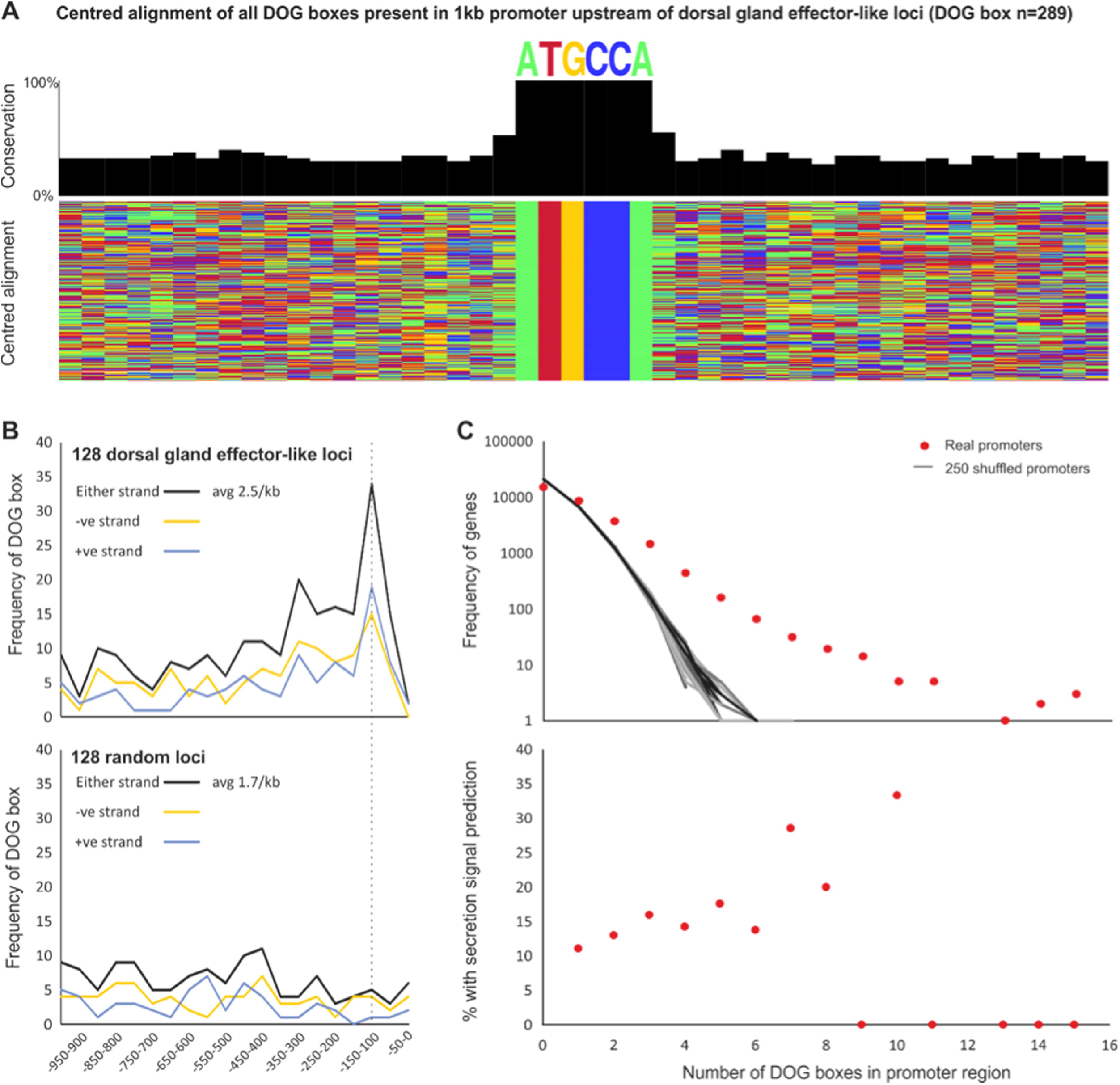

Given that DOG boxes are only present in some effector promoters, to identify a comprehensive repertoire of effectors we combined a number of methods and criteria. First, we aligned the 80 known *H. glycines* effector sequences to the genome using GMAP, identifying 121 putative effector genes, some of which were also genes containing the DOG motifs identified above. Second, the protein motif finder MEME was used on the 80 known effectors, identifying 24 motifs in 60/80 effectors (Figure 3A). One motif (motif 1) was a known signal peptide found in 10/60 effectors [20]. In addition, motifs 8, 12, and 18 were also found at the N terminus in 7/60, 16/60, and 17/60 effectors, respectively. Other genes in the genome that also contain these motifs may also be effectors. Therefore these 24 motifs (Supplemental Data 1) were searched against the proteins predicted in this genome using FIMO, revealing a set of 292 proteins with at least one effector-like motif. Finally, this set of 292 was merged with the 121 putative effector genes and the 160 dorsal effector-like genes mentioned above to produce a unique set of 431 effector-like genes. This gene set was used in downstream analyses exploring effector evolution. Of the 431 effector-like loci, 216 are predicted to encode a secretion signaling peptide and lack a transmembrane domain. While the remaining 215 effector-like loci may contain non-effectors, they were retained for downstream evolutionary analyses because they may represent genes with non-canonical secretion signals, “progenitor” housekeeping genes that gave rise to effectors (e.g. GS-like effectors[19], SPRY-SECs [21], etc.), or an effector graveyard.

To gain further insights into the prevalence of alternative splicing within known effector proteins, the 80 previously identified secreted effector proteins were associated with 371 transcripts. This differs from previous work where 395 transcripts were associated with the 80 effectors in a de-novo transcriptome approach [17]. The main types of alternatively spliced variants for the effector genes included 73 (19.7%) intron retention, 26 (7.0%) alternative 5’ donor site, 25 (6.7%) alternative 3’ acceptor site, 43 (11.6%) alternative transcription start site, 47 (12.7%) alternative transcription termination site, 4 (1.1%) single exon skipping, and 30 (8.1%) multiple exon skipping.

To explore effects that alternative splicing may have on the protein function, functional domain analysis was conducted using the Pfam domain annotation tool [22]. Of the 69/80 single copy effectors genes, only 9 (7.7%) with 51 corresponding isoforms had identifiable functional protein domains. In total, our analysis identified 12 protein functional domains for the 9 effector genes (on average 0.24 domains per an isoform). Each of the 9 effector genes had at least one AS event that altered the predicted domain architecture. Overall, the transcripts included domain architectures with no change, with at least one added, modified, or deleted functional domain.

Horizontal gene transfer (HGT) was important for the evolution of parasitism in the root-knot and cyst nematodes [23–30]. To better understand the role of HGT in the evolution of effectors in *H. glycines*, we calculated an Alien Index (AI) for each transcript using a ratio of similarity to metazoan and non-metazoan sequences [31]. A total of 1,678 putative HGT events (AI>0) were observed in the predicted *H. glycines* proteome (Supplemental Data 2), which are distributed on 461 different contigs (Figure 3B). This prediction includes 151 genes with strong HGT support (AI>30) (Figure 3B), and 82 genes previously identified in closely related nematodes (Table S4). The number of introns was significantly reduced in genes with AI>0 (6.8 vs 9.7, p<0.001, Student’s t-test) (Figure 3B), further supporting their non-metazoan origin. Among these, the highest E-values were of bacterial, fungal, or plant origin for 70.8% (114/161), 19.3%, and 9.9%, respectively (Supplemental Data 2). Interestingly, only 7/151 high confidence HGT genes were co-identified as one of the 431 effector-like loci.

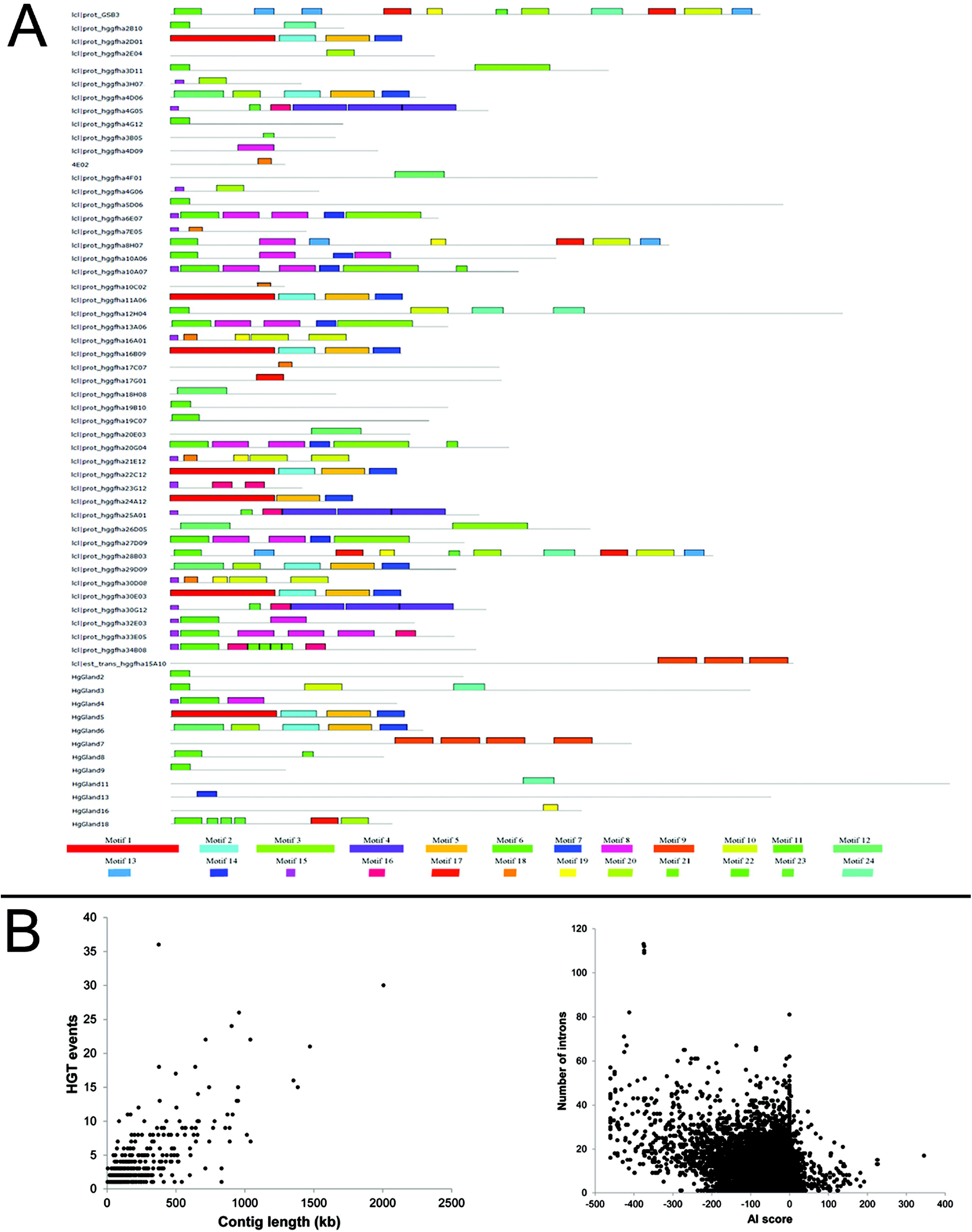

The tandem duplication (TD) of genes in pathogen genomes is a common evolutionary response to the arms race between pathogen and host as a means to avoid/overcome host resistance [32]. To identify genes involved in tandem duplications as a measure to discover sources of virulence genes, we implemented RedTandem to survey the *H. glycines* genome. We determined that a total of 18.7MB of the genome is duplicated with a total of 20,577 duplications in the genome. While most individual duplications were small, the average tandem duplication size was 909bp (Figure 4). We verified that tandem duplications were not assembly artifacts by aligning the PacBio preads to the genome and confirmed that the larger than average tandem duplications (4410/4241) were spanned by PacBio preads across >90% of tandem duplication length. We determined that the density of genes in the tandemly duplicated regions is higher than in non-duplicated regions of the genome: 6,730/18.7MB (~360 genes/MB) vs 23,039/105.2MB (~219 genes/MB), and thus contributes to one fifth of the total gene count in the *H. glycines* genome. The largest groups of orthologous genes found in tandem duplications (881/3940 genes) were annotated with BLAST to the NCBI non-redundant (NR) database, revealing that the 38 largest clusters of duplicated genes were frequently transposable element genes, effector/gland-expressed genes, or BTB/POZ domain-containing genes (Figure S5). Both effector-like loci (136/431; 36%) and HGT genes (38/151; 25%) were duplicated in the tandem duplications. Of effectors that were orthologous in the tandemly duplicated orthologs, Hgg-20 (144), 4D06 (11) and 2D01 (11) were the most frequent, while RAN-binding proteins formed the largest cluster of HGT genes (Figure S5).

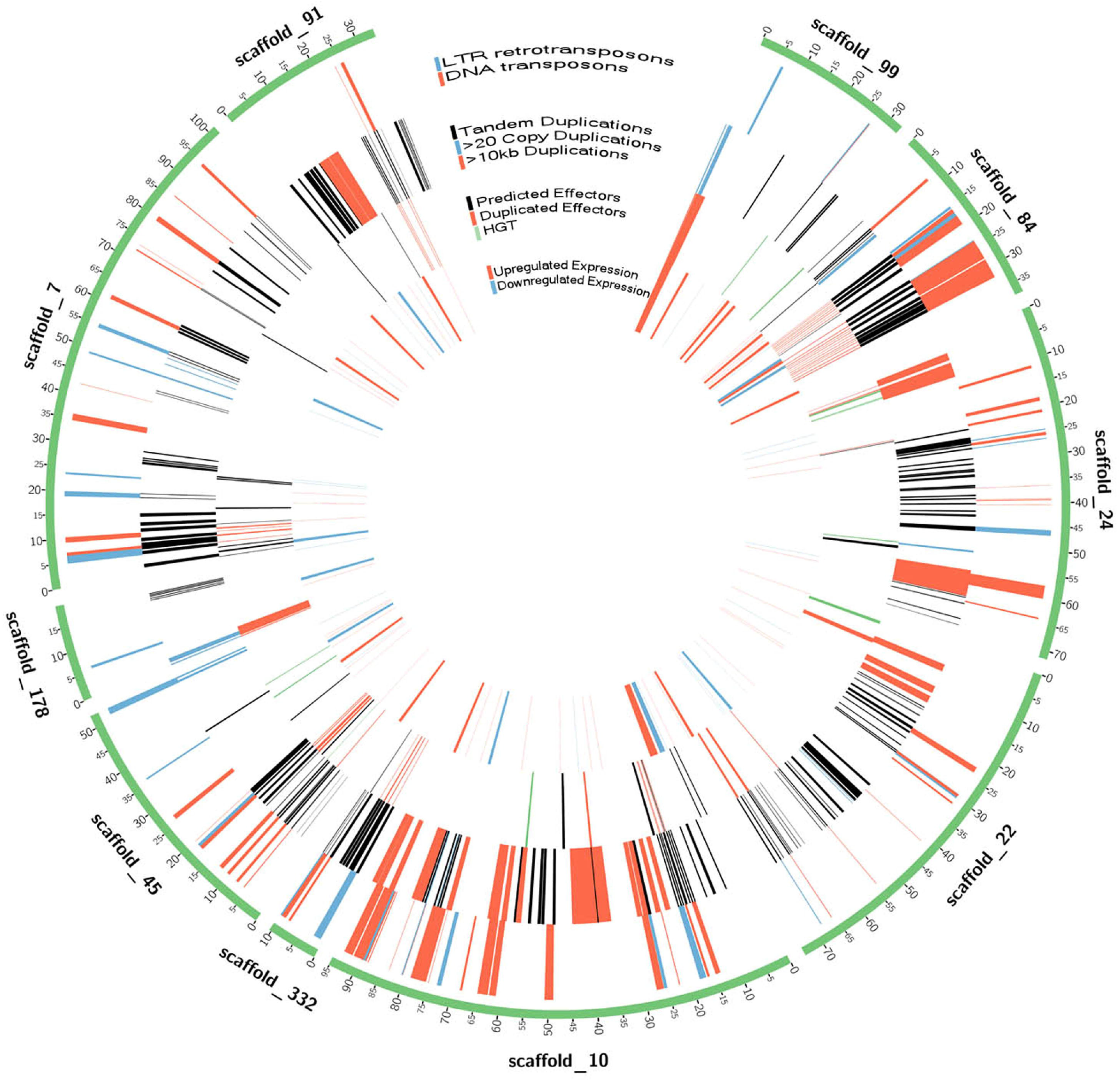

To investigate whether transposons were associated with the expansion of effector genes, we created confident transposon and retrotransposon models using data co-integrated from RepeatModeler, LTR finder, and Inverted Repeat Finder (see methods). One-third of the *H. glycines* genome was considered repetitive by RepeatModeler (32%, 39Mb) with the largest classified types being DNA transposons (7.53%), LTR elements (2.92%), LINEs (1.83%) and SINEs (0.04%) (Table S5). To identify full-length DNA transposons and LTR retrotransposons, Inverted Repeat Finder (3.07) and LTR Finder (1.0.5) were used to identify terminal inverted repeats and LTRs, respectively. The genomic co-localization of RepeatModeler repeats and inverted repeats led to the identification of 1,075 DNA transposons with a mean size of 6.6kb and encompassing 1,915 genes (Figure 4). Similarly, the overlap of RepeatModeler repeats and LTR Finder repeats identified 592 LTR retrotransposons with 8.1kb mean size and encompassing 1,401 genes (Figure 4). Among the genes found within DNA or retro-transposon borders, 58/1,075 and 22/1,401 were effector-like, respectively. However, transposon-mediated duplication is not specific to effectors, as evidenced by 14 duplicated non-effector HGT genes. To obtain a measure of duplications associated with transposons, Bedtools intersect was used to identify transposon-associated gene overlap with tandemly duplicated genes. Of the 6,767 genes contained in tandem duplications, 969 and 656 were contained in DNA and LTR transposons, respectively.

Another possible mechanism by which *H. glycines* could overcome soybean resistance is through changes in coding sequences that result in differences among closely related effectors. Therefore, identifying SNPs in effector genes may reveal mutations associated with effector diversification. Using GATK best practices [33], 1,619,134 SNPs were identified from 15 bulked, pooled DNA preparations from isolate populations of virulent and avirulent *H. glycines* lines. To better understand population-level dynamics SNP-Relate was used to create a PCA plot, and as expected, populations primarily grouped by their original ancestral population but also by selection pressure on resistant cultivars (Figure S6). The SNP density for each gene was determined by dividing SNP frequency by CDS length, and Fisher’s exact tests with the GeneOverlap R package were used to identify significant associations with genes in the 10th and 90th percentile of SNP density (Figure 5). SNP-dense genes were significantly enriched for genes found in tandem duplications, DNA transposons, LTR retrotransposons, and any gene with exon-overlapping repeats. While mutations are present in effectors, effector genes were not associated with high SNP density, although the lack of unique reads in highly duplicated regions may be responsible. Supporting this hypothesis, genes and effectors found in tandem duplications, DNA transposons, and LTR retrotransposons significantly overlapped with the 4,613 genes lacking SNPs, and thus unique sequence reads (Figure 5).

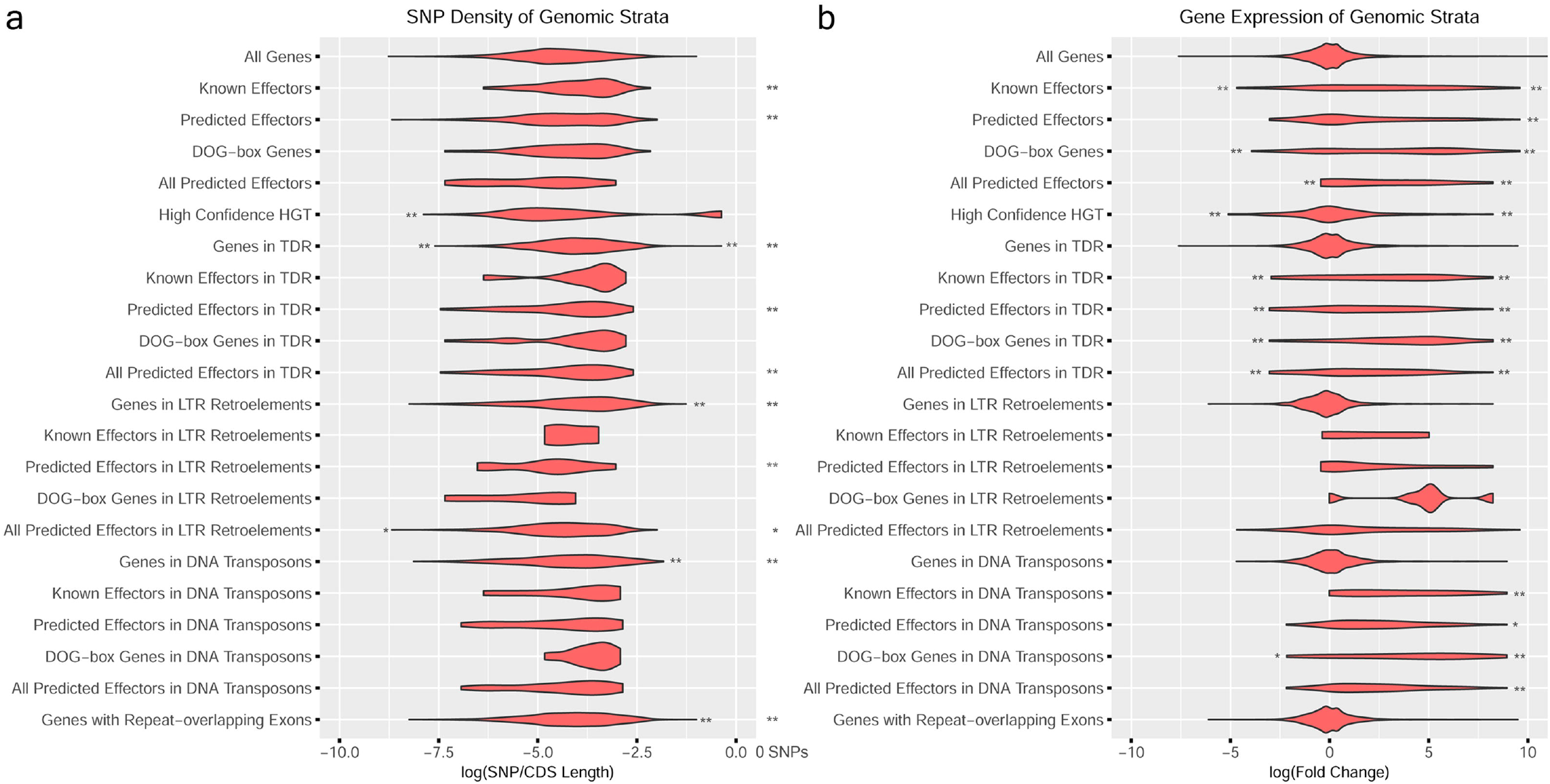

To assess the importance of genes affected by duplications, repeat-association, and SNP density, we utilized gene expression from second-stage juveniles of *H. glycines* population PA3 before and after root infection of a resistant and susceptible soybean cultivar (SRP122521). Genes differentially up and downregulated after infection were identified using DESEQ with a q-value cutoff of 1e-8, revealing 1,211 and 568 genes with significant up and down regulation, respectively. To associate differential expression with effectors and other gene categories, significant associations were identified using the GeneOverlap R package (Figure 5, Table S6). As expected, many of the predicted effectors were significantly upregulated upon infection, a trend that continued with putative effectors found in DNA transposons and tandem duplications. In contrast, the only significantly upregulated gene categories not directly associated with predicted effectors were secreted genes and genes associated with an effector-associated repeat (Family-976 repeat).

However, since virulence genes have a limited span of use before host immunity is developed, the expression of a recognized effector may hinder survival, thus finding effectors with reduced gene expression is not surprising. Generally, genes associated with tandem duplications, HGT, and transposons had similar distributions of expression as genes that were non-associated, yet effectors found in tandem duplications and DNA transposons were significantly enriched for genes with high and low expression (Figure 5). This high and low expression trend in effectors was also apparent in secreted genes at a higher significance, indicating that many potential effectors remain elusive to detection.

## Discussion

To overcome the expected assembly problems associated with high-levels of repetitive DNA and to reveal the evolutionary means behind the rapid evolution and population shifts in *H. glycines*, we used long-read technology to assemble a genome from a heterogenous population of individuals. A number of analyses confirmed a high level of genome completeness with ~88% of the RNA-Seq aligning, 93% of preads aligning, and zero contaminating scaffolds (Table S1, Figure S1). While percentages of missing BUSCO [34]genes were high, BUSCO genes were 72% complete, ranking *H. glycines* the best among sequenced genomes in the cyst and knot-nematode clades (Figure 1, Table S2). Some level of artifactual duplication may be present in the genome, with BUSCO gene duplication being highest among the species analyzed. However, only 79/349 duplicated BUSCO genes are found in tandem duplications, indicating that duplication or heterozygous contigs may be present elsewhere in the genome. With a goal-oriented approach of capturing all genic variation in the genome, we sequenced a population of multiple individuals. We therefore assembled a chimera of individuals, with some duplicated genes originating from single variants in the population. However, even when considering that nearly nine thousand genes could be attributed to repetitive elements and tandem duplications, the gene frequency (20,830) and exon statistics of *H. glycines* are elevated in relation to sister Tylenchida species.

Using hundreds of thousands of individuals, we shed light on evolutionary underpinnings of virulence and parasitism in an *H. glycines* pan-genome. Because plant-parasitic nematodes have many well-documented cases of HGT [35], we investigated the potential role HGT may have in *H. glycines*. Almost all previously identified HGT in plant-parasitic nematodes were also found in *H. glycines* (n=84) (Table S4) [36]. Genes with strong AI (>30) were mainly hydrolases, transferases, oxidoreductases or transporters (Data S2). Of particular interest were genes originating from bacteria or fungi, but lacking BLAST hits to Metazoan species (highlighted blue in Data S3). Among these is a gene coding for an Inosine-uridine preferring nucleoside hydrolase (Hetgly.000009703; AI = 101.2), an enzyme essential for parasitism in many plant-pathogenic bacteria and trypanosomes [37]. A candidate oomycete RxLR effector [38] was also identified in the genome (Hetgly.000002962, Hetgly.000002964 and Hetgly.000002966; AI up to 42.2). Besides being necessary for successful infection, RxLR effectors are also avirulence genes in some species, including the soybean pathogen *Phytophthora sojae* [39]. The *H. glycines* genome is also host to a putative HGT gene (Hetgly.000001822 and Hetgly.000022293; AI up to 55.3) that has been characterized as *a G. pallida* effector (Gp-FAR-1) involved in plant defense evasion by binding plant defense compounds [40]. Thus, horizontal gene transfer appears to contribute to the evolution of *H. glycines* virulence as well as to the ancestral development of parasitism in plant-parasitic nematodes [35, 41–43].

Although HGT is more common among nematodes and arthropods than other animals [44], there are many documented cases of gene duplication leading to evolutionary novelty and phenotypic adaptation across metazoans [45, 46]. With over a fifth of the genes in the *H. glycines* genome found in tandem duplications, characterizing the largest clusters of orthologous gene families in tandem duplications provides relevant information for identifying genes related to parasitism, adaptation, and virulence. A functional assessment of the 38 largest clusters of tandemly duplicated orthologues were largely transposon-associated proteins or proteins related to effectors, indicating that transposons have a role in duplicating effector genes (Figure S5). Because many of the LTRs and TIRs were nested, effector exon shuffling could also be attributed to the frequent rearrangements of nested clusters of transposons, as has been seen in other systems [47]. Genes within the tandemly duplicated regions are also potentially subject to higher rates of mutation, indicated by the significant associations of genes in duplicated regions and SNP density across populations. Yet transposable element-associated effectors were not always silenced, in fact these effectors were some of the most highly upregulated and downregulated genes upon infection (Figure 5).

By merging genomics and transcriptomics data, we can provide important insights into the molecular mechanisms regulated through alternative splicing. Alternative splicing is highly prevalent within *H. glycines*, echoing similar observations in other nematode species [48]. Approximately 70% of the alternative splicing events significantly alter the length of the open reading frame (ORF). The most prevalent alternative splicing event type in *H. glycines* transcriptome is intron retention (30%). Considering the known effector genes, alternative splicing is involved across both the juvenile and adult stages as well as during the infection of either a susceptible or resistant host. The most prominent type of alteratively spliced variant within the known effector genes was intron retention (19.7%), a phenomenon previously identified in the chorismate mutase effector of *G. rostochiensis[49]*. Alternative splicing is often found to alter the protein domain architecture of the effector genes, where the protein domains are deleted or modified.

## Conclusions

The *H. glycines* genome assembly and annotation has provided a glimpse into host and parasite interplay by characterizing known and predicted effector genes, investigating mechanisms of gene birth, and identifying horizontally acquired genes. Further investigation will reveal how effector genes are hitchhiking with transposable elements and the functional relevance of this association. Through extensive characterization of the *H. glycines* genome, we provide new insights into *H. glycines* biology and shed light onto the mystery underlying these complex host-parasite interactions. This genome sequence is an important prerequisite to enable work towards generating new resistance or control measures against *H. glycines*.

## Methods

### Nematode culture and DNA/RNA isolation

*H. glycines* inbred population TN10, Hg type 1.2.6.7, was grown on susceptible soybean cultivar Williams 82 in a greenhouse at Iowa State University. A starting culture of approximately 10,000 eggs from Dr. Kris Lambert, University of Illinois, was bulked for four generations on Williams 82 soybeans grown in a 2:1 mixture of steam pasteurized sand:field soil in 8” clay pots, with approximately 16h daylight at 27□. Genomic DNA was extracted from approximately 100,000 eggs in a subset of third generation cysts. Egg extraction was performed with standard nematological protocols [50], eggs washed 3 times in sterile 10 mM MES buffered water, and pelleted before flash freezing in liquid nitrogen.

Genomic DNA was isolated using the MasterPure Complete DNA Purification Kit (Epicentre) with the following modifications: Frozen nematode eggs were resuspended in 300 ul of tissue and cell lysis solution, and immediately placed in a small precooled mortar, where the nematode solution refroze and was finely ground. The mortar was then placed in a 50□ water bath for 30 minutes, then transferred to 500 ul PCR tubes with 1 ul of proteinase K, and incubated at 65□ for 15 minutes, inverting every 5 minutes. Genomic DNA was resuspended in 30 ul of RNAse/DNase free water, quantified via nanodrop, and inspected with an 0.8% agarose gel at 40V for 1h. Two 20 kb insert libraries were generated and sequenced on 20 PacBio flow cells at the National Center for Genome Resources in Santa Fe, NM (SRR5397387 – SRR5397406).

Fifteen *H. glycines* populations were chosen based on Hg-type diversity and were biotyped to ensure identity (TN22, TN8, TN7, TN15, TN1, TN21, TN19, LY1, OP50, OP20, OP25, TN16, PA3, G3). Genomic DNA from approximately 100,000 eggs for each population was extracted as described previously, and 500 bp libraries were sequenced on an Illumina HiSeq 2500 at 100PE (SRR5422809 – SRR5422824)

Six life stages were isolated for both PA3 and TN19 *H. glycines* populations: eggs, pre-parasitic second-stage juveniles (J2), parasitic J2, third-stage juveniles (J3), fourth stage juveniles (J4) and adult females. Parasitic J2 were isolated, followed by isolations of J3, J4, and adult females at 3, 8, 15, and 24 days post-infection via a combination of root maceration, sieving and sucrose floatation, using standard nematological methods[50]. Total RNA was extracted with the Exiqon miRCURY RNA Isolation Kit (Catalog #300112). RNA was combined to form three pools for each population, corresponding to early (egg and pre-parasitic J2), middle (parasitic J2 and J3) and late (J4 females and early adult females) developmental stages. The IsoSeq data were used to improve the annotation (see below) (SAMN08541516-SAMN08541521).

### Genome Assembly

A PacBio subreads assembly was generated with Falcon to correct subreads into consensus preads (error corrected reads), followed by contig assembly. An alternative approach using only transcript containing preads was helpful in solving heterozygosity and population problems. Transcripts were aligned to preads using Gmap [51], and a pool of preads for each unique transcript was assembled using CAP3 [52]. The longest assembled contigs and all unassembled preads were retained and read/contig redundancy was removed with sort and uniq. New FASTA headers were generated using nanocorrect-preprocess.pl [https://github.com/jts/nanocorrect/blob/master/nanocorrect-preprocess.pl], and sequences were then assembled with Falcon into 2,692 contigs (supp file *H. glycines.cfg)*. Falcon output was converted to Fastg with Falcon2Fastg [https://github.com/md5sam/Falcon2Fastg.py], and longer scaffolds were created with Bandage [53] using multiple criteria. 1) The longest path was chosen and ended with an absence of edges. 2) If the orientation of an interior contig was disputed, one set of edges was deleted to extend the scaffold. 3) The shortest path through difficult repetitive subgraphs was chosen.

Intragenomic synteny was used to remove clonal haplotigs [54, 55] (synteny as below). When synteny was identified between two contigs/scaffolds, if a longer 3’ or 5’ fragment could be made, then the ends of each contig/scaffold were exchanged at the syntenic/nonsyntenic juncture. All remaining duplicate scaffolds retaining synteny were truncated or removed from the assembly, and followed by a BWA [56] self-alignment to remove redundant repetitive scaffolds.

### Genome Quality Control

Multiple measures were taken to assess genome assembly quality, including a BLASR [57] alignment of PacBio subreads, preads, and ccsreads resulting in alignment percentages at 88.7, 93.3, 90.1%, respectively (Table S1). Gmap and Hisat2 (2.0.3) mapped 86.4% percent of a transcriptome assembly and ~88% of the five RNA-seq libraries, respectively (Table S1). Genome completeness was assessed with BUSCO [34] at 71.9 %. An absence of contamination was found with Blobtools (4.8.2) [58] using MegaBlast (2.2.30+) to the NCBI nt database, accessed 02/02/17, at a 1-e5 e-value. See supplementary methods for more detail.

### Genome Annotation

To account for the high proportion of noncanonical splicing in nematodes [12], Braker [59] was used to predict genes using Hisat2 (2.0.3) [60] raw RNA-Seq alignments of ~23 million 100bp PE RNA-Seq reads and GMAP [61] alignments of IsoSeq reads and all EST sequences from NCBI. Because gene models were greatly influenced by repeat masking, three differentially repeat-masked genomes were used for gene prediction: unmasked, all masked, and all except simple repeats masked (see supp table RNASEQ mapping in excel). All protein isoforms were annotated with Interproscan [62, 63] in BlastGO [64], and with BLAST [65] to Swiss-prot [66] and Uniref [67].

### Repeat Prediction

Repetitive elements in the *H. glycines* genome were classified into families with five rounds of RepeatModeler (1.0.8) [68], followed by genome masking with RepeatMasker [69]. Inverted Repeat Finder (3.07) and LTR Finder (1.0.5) were used to define the border of a TE only when overlapping Repeatmodeler repeats were present. Supplemental helitron prediction was done with Helitronscanner [70].

### Promoter analyses

To determine to what extent cyst nematodes use common mechanisms for dorsal gland effector regulation, a robust list of putative effectors was collated. The *G. rostochiensis* DOG-effectors [12] were used as query in blastp to identify DOG-effector-like loci in the predicted proteome of *H. glycines*. The most similar sequence was retrieved if it was identified with an evalue <1e-10 and it encoded a putatively secreted protein (78 unique *H. glycines* loci). Using the same approach, sequences similar to other published dorsal gland expressed effectors were identified [6, 71] and combined with the DOG-effector-like list to a non-redundant 128 loci. A 500 bp region 5’ of the ATG start codon, termed the promoter region, was extracted from these 128 loci and used for motif enrichment analysis using HOMER [72], as previously described [12]. DOG-box positional enrichment was calculated using FIMO web server [73] and predictive power calculated using custom python scripts.

### Effector Analyses

Gmap [61] was used to align 80 previously identified effectors to the genome [6, 71, 74, 75]. Conserved protein motifs in effectors were identified with MEME: -nmotifs 24, -minsites 5, -minw 7, -maxw 300, and zoops (zero or one per sequence) [76]. These motifs were used as FIMO queries to search the inferred *H. glycines* proteome [76].

### Synteny

The genome, gff, and peptide sequences for *C. elegans* (WBcel235), *G. pallida* [77], and *M. hapla* [78] were downloaded from WormBase [79]. The genome and gff of *G. rostochiensis* [12] was downloaded from NCBI. The *G. ellingtonae* genome was also downloaded from NCBI[80], but gene models were unavailable, thus gene models for *G. ellingtonae* were called with Braker using RNA-seq reads from SRR3162514, as described earlier.

Fastp and global alignments with Opscan (0.1) [81] were used to calculate orthologous gene families between *H. glycines* and *C. elegans [82], G. pallida [77], G. ellingtonae [80], G. rostochiensis [12], M. hapla [83]*, and *M. incognita [14]*. All alternatively spliced variants and all possible multi-family genes were considered.

To infer synteny, iAdHoRe 3.0.01 [84] was used with prob-cutoff=0.001, level 2 multiplicons only, gap-size=15, cluster-gap=20, q-value=0.9, and a minimum of 3 anchor points. Syntenic regions are displayed using Circos (0.69.2) [85].

### Phylogenetic Tree

Predicted protein sequences from the aforementioned nematode genomes (excluding *C. elegans*) were scanned with BUSCO 2.0 [34] for 982 proteins conserved in *nematoda*. 651 proteins were found in at least 3 species and aligned with Prank [86] in Guidance [87]. Maximum likelihood gene trees were computed using RAxML [88] with 1000 bootstraps and PROTGAMMAAUTO for model selection. Astral [89] was used to prepare a coalescent-based species tree.

### Tandem duplication

ReDtandem.pl was used to identify tandem duplications in the genome [90]. Tandem duplicate orthologous genes were identified using a self-BlastP to predicted proteins with 50% query length and 90% identity [65]. To annotate clusters of orthologous genes, groups of highly connected nodes or entire clusters were concatenated and queried with BlastP to the NCBI NR database [91].

### SNP density and PCA analysis of fifteen *H. glycines* populations

Raw sequences from fifteen populations of *H. glycines* nematodes were quality checked with FastQC [92]. Reads were aligned to the *H. glycines* genome using BWA-MEM [56]. The BAM files were sorted, cleaned, marked for duplicates, read groups were added and SNP/Indel realignment were performed prior to calling SNPs and Indels with GATK. Custom Bash scripts were used to convert the vcf file into a gff for use with Bedtools (2.2.6) to identify SNP and exon overlap [93]. The density of SNPs was calculated by dividing the number of SNPs/CDS length (bp). Phasing and imputing SNPs with Beagle 4.1 [94, 95] followed by a PCA analysis of SNPs vs Hg-type virulence using SNPRelate (1.12.2) [96].

### RNA-seq expression

RNA-seq reads were obtained from NCBI SRA accession SRP122521. Briefly, SCN inbred population PA3 was grown on soybean cultivar Williams 82 or EXF63. Pre-parasitic second-stage juveniles and parasitic second stage juveniles were isolated from roots of resistant and susceptible cultivars at 5 days post-inoculation [17]. 100bp PE reads were aligned to the genome using HiSat2 [60]. Read counts were calculated using FeatureCounts from the Subread package [97], followed by Deseq2 [98] to determine log-fold change between the pre-parasitic samples (2 x ppJ2_PA3) and parasitic J2 samples (2 x pJ2_s63, pJ2_race3_Forrest).

### Alternative splicing analysis

The analysis of the global changes and effector specific effects in alternative splicing landscape was assessed following a recent *de novo* transcriptomics analysis of the *H. glycines* nematode effectors [17] Transcriptome annotation was constructed using 230 million RNA-Seq reads from both pre-parasitic and parasitic J2 *H. glycines* [17], 34,041 iso-seq reads from three life stages of both a virulent and an avirulent strain, and H. *glycines* ESTs in NCBI (35,796). Specifically, using a standard alternative splicing analysis pipeline [99], 230 million reads from both pre-parasitic and parasitic J2 *H. glycines* [17] were preprocessed with Trimmomatic [100], aligned with Tophat 2.1.1 [101], and quantified with Cufflinks 2.2.1 [102], followed by conversion of FPKM to TPM [103], and patterns assessment with IsoformSwitchAnalyzerR [104]. For the 80 previously identified effectors [6, 71, 74, 75], the changes in the functional domain architectures between specific alternatively spliced isoforms are determined using InterPro domain annotation server with a focus on Pfam domains [105].

### Bioinformatics scripts

Scripts used for the alternative splicing analysis can be found at https://github.com/bioinfonerd/SCN_AS_RNA_Seq. Scripts used for the promoter analysis can be found here: https://github.com/sebastianevda/Fimo_parse/tree/master. All other scripts and bioinformatic analyses can be found at: https://github.com/ISUgenomics/SCNgenomepaper/tree/master/SCNgenome/Camtech738GenomeAnalyses.

## Declarations

### Ethics approval and consent to participate

Not applicable

### Consent for publication

Not applicable

### Availability of data and materials

Datasets generated during the current study are available at Genbank accessions (SRR5397387 – SRR5397406), (SRR5422809 – SRR5422824), (SAMN08541516-SAMN08541521).

### Authors’ contributions

RM, TRM, PSJ, MGM, MH, AJS, and TJB conceived and designed the experiment. TRM isolated and acquired the data. RM and AJS performed the assembly. SEvdA performed and wrote the promoter analysis. DK and NTJ performed and wrote the alternative splicing analysis. BM and EL performed and wrote the horizontal gene transfer analysis. RM performed all other comparative analyses. All authors made substantial contributions to the final text.

